# From Data to Insights: Machine Learning Empowers Prognostic Biomarker Prediction in Autism

**DOI:** 10.1101/2023.11.13.566549

**Authors:** Ecmel Mehmetbeyoglu, Abdulkerim Duman, Serpil Taheri, Yusuf Ozkul, Minoo Rassaulzadegan

**Affiliations:** Department of Cancer and Genetics, Cardiff University, Cardiff, United Kingdom; Betul-Ziya Eren Genome and Stem Cell Center, Erciyes University, Kayseri, Turkey; School of Engineering, Cardiff University, Cardiff, United Kingdom; Department of Medical Biology, Erciyes University, Kayseri, Turkey; Department of Medical Genetics, Erciyes University, Kayseri, Turkey; Université Côte d’Azur, CNRS, Inserm, France

**Keywords:** Autism, miR-126-3p, Machine Learning

## Abstract

Autism Spectrum Disorder (ASD) poses significant challenges to society and science due to its impact on communication, social interaction, and repetitive behaviour patterns in affected children. The Autism and Developmental Disabilities Monitoring (ADDM) Network continuously monitors ASD prevalence and characteristics. In 2020, ASD prevalence was estimated at one in 36 children, with higher rates than previous estimates. This study focuses on ongoing ASD research conducted by Erciyes University. Serum samples from 45 ASD patients and 21 unrelated control participants were analysed to assess the expression of 372 microRNAs (miRNAs). Six miRNAs (miR-19a-3p, miR-361-5p, miR-3613-3p, miR-150-5p, miR-126-3p, and miR-499a-5p) exhibited significant downregulation in all ASD patients compared to healthy controls. The current study endeavours to identify dependable diagnostic biomarkers for ASD, addressing the pressing need for non-invasive, accurate, and cost-effective diagnostic tools, as current methods are subjective and time intensive. A pivotal discovery in this study is the potential diagnostic value of miR-126-3p, offering the promise of earlier and more accurate ASD diagnoses, potentially leading to improved intervention outcomes. Leveraging machine learning, such as the K-nearest neighbours (KNN) model, presents a promising avenue for precise ASD diagnosis using miRNA biomarkers.

## 1. Introduction

Autism Spectrum Disorder (ASD) presents a challenge both for society and science, as it affects a high number of children with alterations in communication, social interaction, and behaviour with repetitive patterns^1–4^. The Autism and Developmental Disabilities Monitoring (ADDM) Network performs continuous monitoring of ASD. It tracks the occurrence and traits of ASD in children who are 8 years old, and whose parents or guardians reside in 11 ADDM Network locations throughout the United States. In 2020, the estimated prevalence of ASD in 8-year-old children was one in 36, which amounts to approximately 4% of boys and 1% of girls. These estimates are higher than the previous ADDM Network estimates from 2018 to 2020^5,6^.

It impacts the most intricate brain functions, prompting important inquiries from neurobiologists, social scientists, and geneticists. The high correlation between monozygotic twins suggests a genetic cause, but the wide range of phenotypes, even among twins, suggests that epigenetic mechanisms play a significant role^7–9^. In the future, the precision in the diagnosis of characters and comprehension of ASD will need to be included in the molecular study of brain functions, which is the highest level of biological complexity. However, in the short term, identifying the cause of ASD could result in beneficial medical advancements, expert advice for both medical professionals and parents, and most importantly, early detection and support for affected children to promote their optimal development. This is crucial for physicians responsible for their care.

Scientific and medical institutions worldwide are currently studying large to extremely large groups of patients, with one recent study including 35,584 individuals, including 11,986 with ASD, their families, and controls ^10,11^. Various genetic research avenues are currently being explored using cutting-edge genome analysis techniques. More than 100 mutated loci with protein-coding alterations were identified in the genomes of patients with autism. Moreover, the detection of tandem DNA repeats through genome-wide analysis has indicated their expansion in individuals with autism, with many expressed in early developmental stages in neurons and neuronal precursors ^12^. Recently, a study reported the role of RNA in controls of CGG repeats in FXS cells with the possibility of FMR1 reactivation in patients’ cells^13^. Uncovering the mechanisms, extent, variability, and connection of the disorder with other neurodevelopmental abnormalities is a high priority. However, none of the genetic variations found were present in all or even most patients, thus disqualifying them as primary determinants of the disorder.

The recent publication from Erciyes University pertains to different areas of ongoing ASD research. The initial phase involved creating a group of 45 patients from 37 separate families, with each family having genetically related individuals (parents, siblings) and 21 unrelated control participants, resulting in a total of 187 serum samples ^14^. The study then concentrated on the group of genes that encode miRNAs, which are 22nt-long non-coding regulatory RNAs recognized as significant factors in cellular differentiation processes^15^. The Kayseri cohort study analysed the expression of 372 miRNAs using miRNA PCR Array profiles and found that six miRNAs (miR-19a-3p, miR-361-5p, miR-3613-3p, miR-150-5p, miR-126-3p, and miR-499a-5p) were significantly down-regulated in the serum of all 45 ASD patients, compared to healthy controls. The fact that all patients showed the same profile of reduced expression of these six miRNAs indicates that this feature is a significant intrinsic characteristic of ASD. The downregulation of these six miRNAs raises a question, can miRNAs be a reliable biomarker in the clinic as a computer-aided diagnostic model?

Artificial intelligence (AI) can be useful for big data with different parameter interpretation processes^16^. The theories and methods used to create automated machines that can mimic human intelligence are collectively referred to as AI. Recent advances in technology have ushered in a new era of precision medicine that combines machine learning (ML) and biological science to analyse diseases using big data. AI includes deep learning, neural networks, and machine learning. Thanks to these strategies, computers are now able to come up with wise selections. ML is commonly employed in the study of cancer and is gaining popularity in the identification and treatment of cancer^17^.

The overall goal of ML is to build automated tools that can quickly classify from previously observed examples and generate by designing or learning functional dependencies between selected input (features) and output (classes) selected^18^. Therefore, the diagnosis of an individual with multi-factorial diseases such as ASD, which aims to translate the knowledge of the characteristics extracted from the miRNA features into meaningful groups (groups of healthy individuals or with ASD), is fundamentally an ML problem.

The purpose of the study is to assess whether ML algorithms, trained on a comprehensive set of miRNA findings collected in advance, can differentiate between healthy individuals and those with ASD and predict the likelihood of ASD in individual patients with accuracy.

MiRNA data that we have generated (q-RT-PCR) has the potential to be a valuable tool for diagnosing ASD and other medical conditions. The hypothesis is that the ML algorithms can successfully identify patterns in miRNA data that distinguish healthy individuals from those with ASD, resulting in high accuracy in predicting the disease in testing data. Therefore, the study aims to investigate the performance of the ML medels in analysing miRNA data and evaluate its potential for clinical application. Our results show KNN models, in particular, showcased outstanding accuracy and overall performance, indicating their potential utility in practical applications within the domain of interest.

## 2. Results

### 2.1. Subject characteristics

Peripheral blood samples were collected from 45 patients with ASD and 37 normal healthy volunteers, the clinical characteristics of which are published in Ozkul et. al 2013 and Rassoulzadegan et. all 2022^14,19^. In the context of a cohort study, the study encompassed the quantification of 372 miRNAs’ expression levels. After the implementation of the Benjamin Hochberg method, it became evident that 180 miRNAs displayed statistically significant disparities among the study groups. To enhance the precision of the selection process, a predefined threshold was introduced to identify miRNAs that exhibited downregulation by at least 90% in comparison to the control group’s expression levels. Employing this stringent selection criterion, the study identified merely six miRNAs that exhibited a statistically significant downregulation.

Figure 1 shows the expression levels of 6 miRNAs in the plasma that were prospectively measured. Among these 6 miRNAs including miR-3613-3p, miR-150-5p, miR-19a-3p, miR-361-5p, miR-499a-5p and miR-126-3p were significantly decreased in patients with ASD and their family.

**Figure 1.**
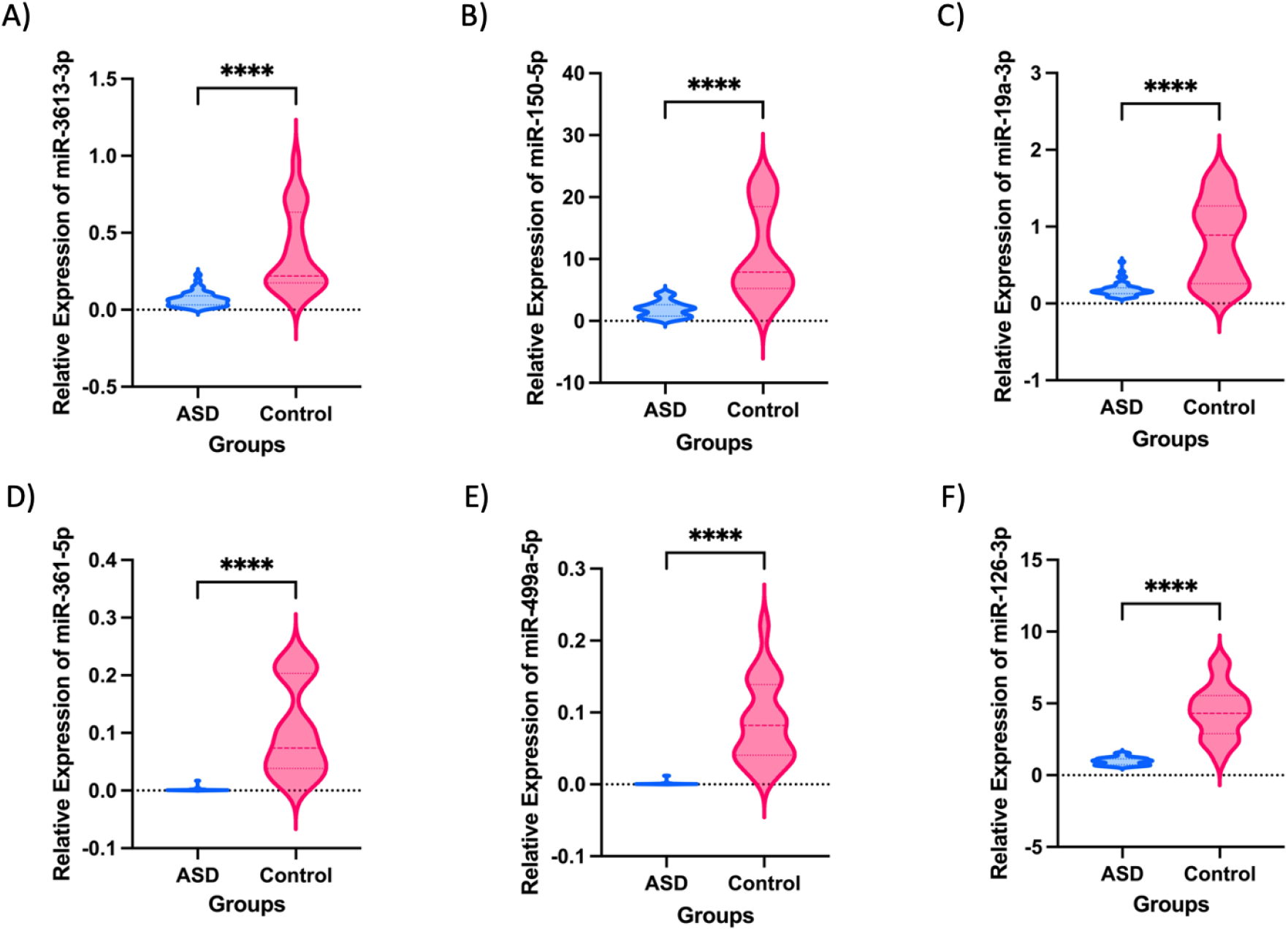
Box violin plots of relative miR-3613-3p (A), miR-150-5p (B), miR-19a-3p (C), miR-361-5p (D), miR-499a-5p (E), miR-126-3p (F) expression level in ASD (n=45) and healthy control (n=21). Asterisks denote significant differences by unpaired t-test, ****<0.0001

### 2.2. ROC analysis

We used data produced by Ozkul et. al 2020 to assess the reliability and performance of the ML program. The study evaluated the diagnostic potential of differentially expressed miRNAs in plasma as biomarkers for the diagnosis of ASD using a receiver operating characteristics (ROC) ML algorithm. As depicted in Figure 2, we implemented the flowchart as our approach for the identification of miRNAs using a machine learning method. The ROC curves of individual miRNAs showed area under curves (AUC) scores ranging from 0.894 to 0.998, indicating good discriminatory power (Figure 3 and Table 1). The miRNA with the highest AUC score, miR-361-5p, was found to have greater potential in discriminating between ASD patients and the control group compared to other miRNAs. A clear separation of six individual miRNA between ASD and control groups appears with AUC values with a 95% confidence interval. The miR-361-5p is the most reproducible diagnostic biomarker for ASD in serum based on ROC curves.

**Figure 2.**
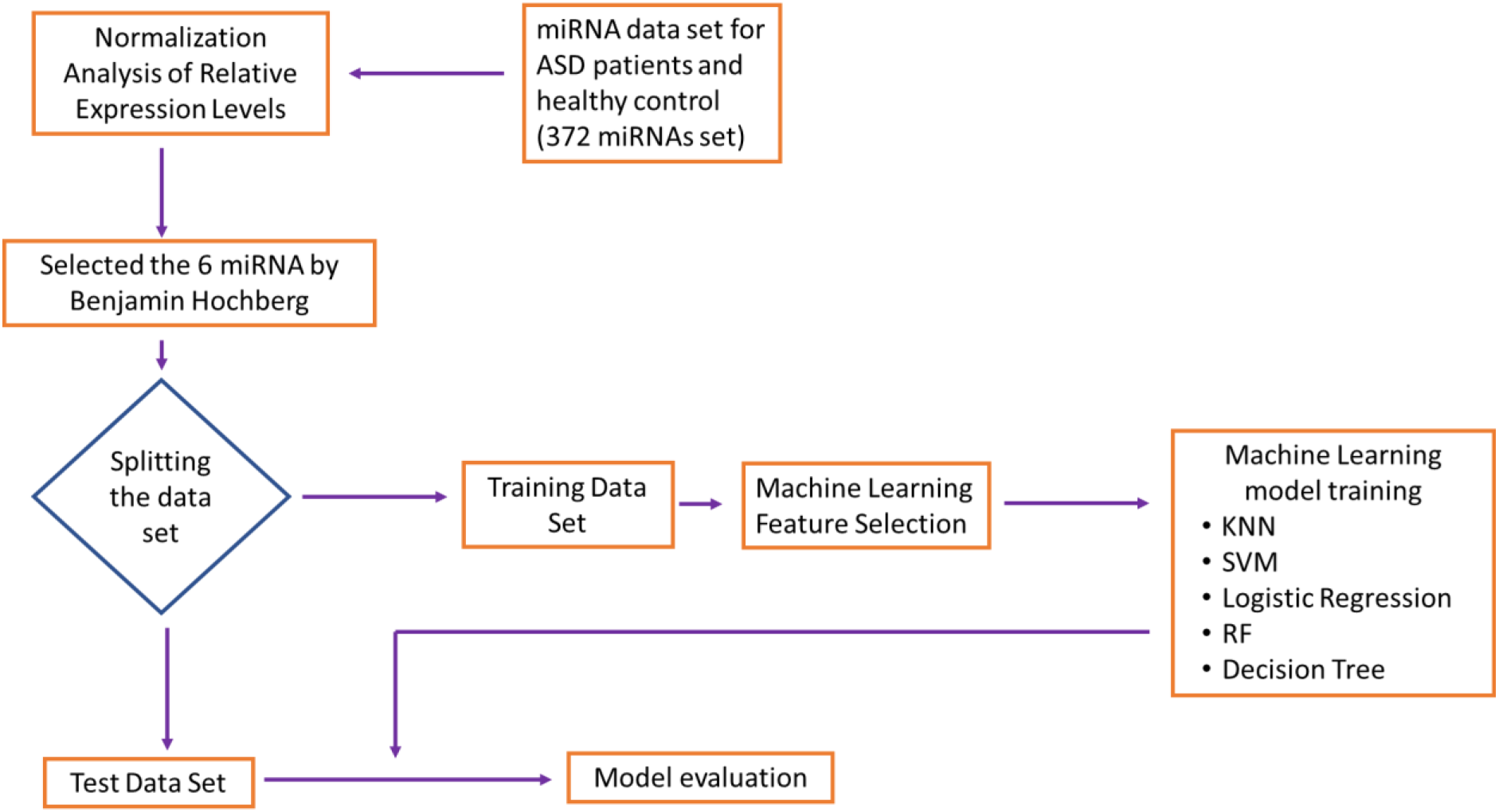
The flowchart of the machine learning method for identification of miRNAs.

**Figure 3.**
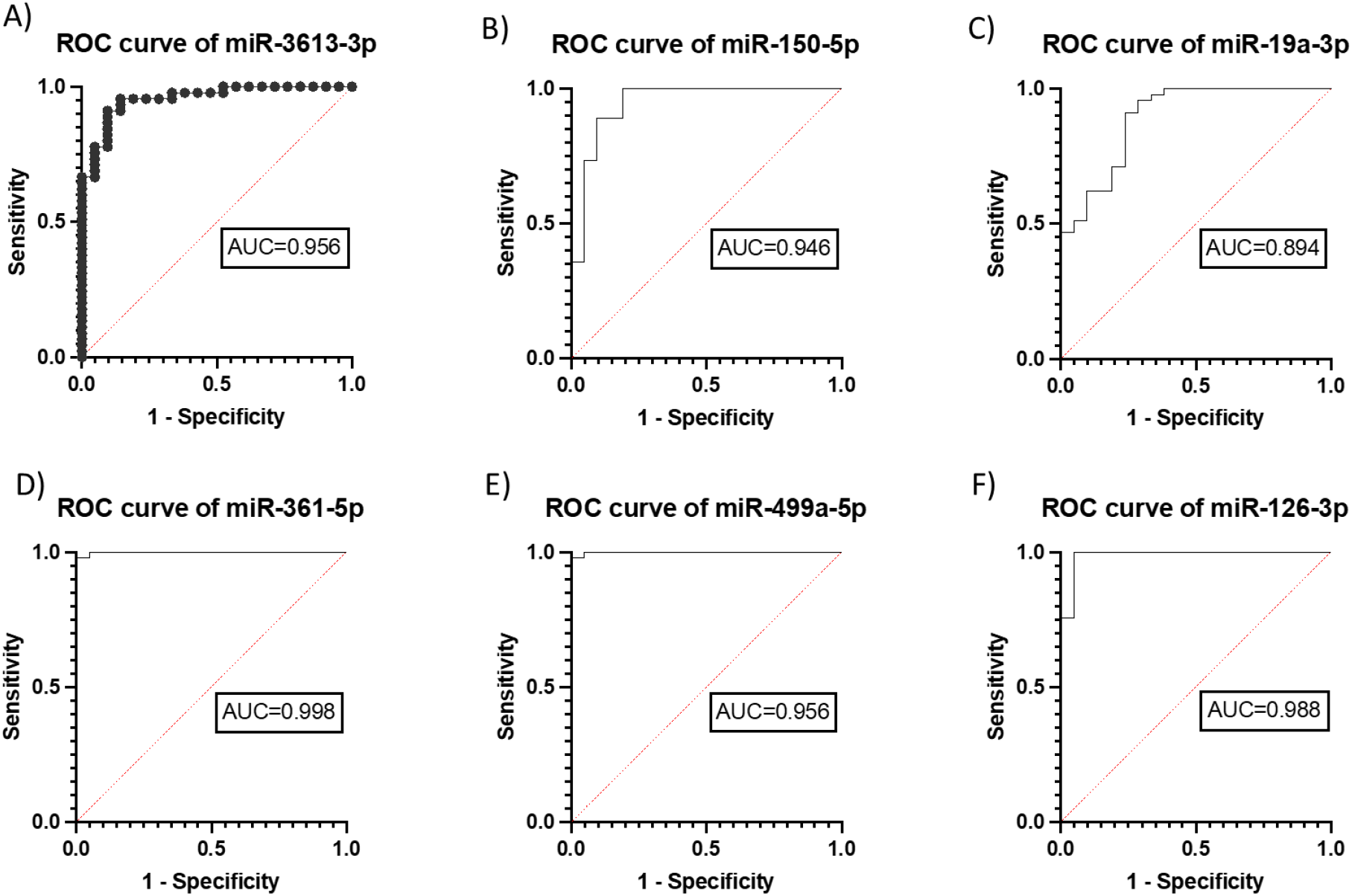
Evaluation of the diagnostic effectiveness of autism-specific miRNAs biomarkers. ROC curve of (A), miR-150-5p (B), miR-19a-3p (C), miR-361-5p (D), miR-499a-5p, miR499a-5p (E), miR-126-3p (F) with an area under the curve (AUC) value. ROC, receiver operating characteristics

**Table 1.** ROC analysis of miRNAs. AUC: Area under the curve, CI: Confidence Interval, SEM: Standard error of the mean.

| miRNAs | AUC | SEM | 95% CI | *p* |
| --- | --- | --- | --- | --- |
| miR-3613-3p | 0.956 | 0.023 | 0.9109-1 | <0.0001 |
| miR-150-5p | 0.946 | 0.033 | 0.8803-1 | <0.0001 |
| miR-19a-3p | 0.894 | 0.043 | 0.8094-0.979 | <0.0001 |
| miR-361-5p | 0.998 | 0.001 | 0.9954-1 | <0.0001 |
| miR-499a-5p | 0.956 | 0.001 | 0.9954-1 | <0.0001 |
| miR-126-3p | 0.988 | 0.0123 | 0.9646-1 | <0.0001 |

### 2.3. Machine Learning

Following getting data, the future selection was applied to data in which relevant features or variables are selected from the pre-processed data. This is an important step that can greatly affect the accuracy of the ML model. In this study, the six significantly expressed miRNAs were selected as features for the ML models based on previous study^14^.

Pearson correlation analysis was conducted on our dataset, leading to the simplification of the model and the reduction of overfitting through the removal of highly correlated features. This enhancement not only enhances model interpretability but also refines its overall performance. Specifically, we focused on the six miRNAs that were previously identified as potential biomarkers for ASD diagnosis. Enrichment scores for 4 of 6 miRNAs had significant Pearson correlations and there were significant differences between patients and controls (Welch’s t-test, p<0.05) (Figure 4). Pearson’s r score was between +0.80 and 1, indicating that the module has a high level of predictive power. These results support the potential utility of these miRNAs as biomarkers for ASD diagnosis and disease activity assessment.

**Figure 4.**
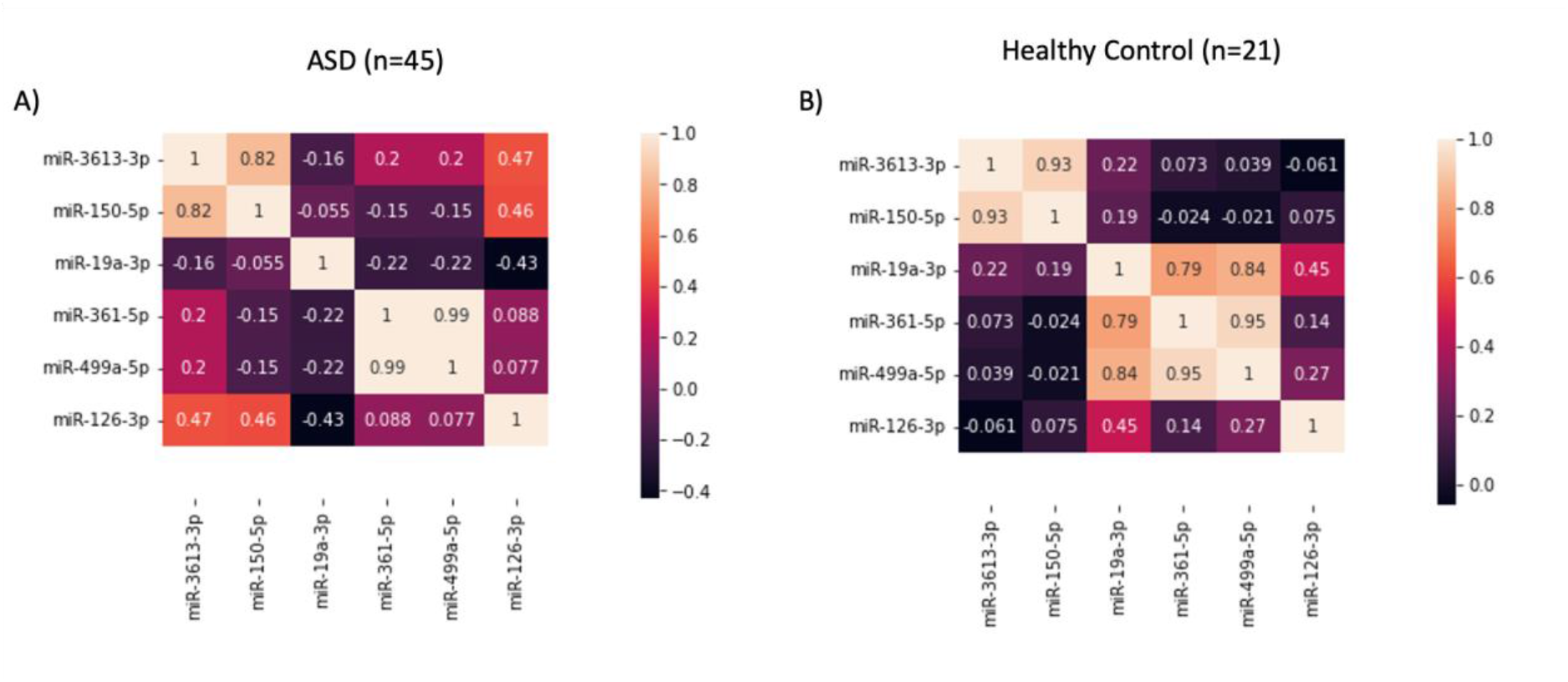
Pearson correlation analysis between six miRNAs. A) Correlations between miRNAs in the ASD patients B) Correlations between six miRNAs in the healthy control.

We found that miR-499a-5p and miR-361-5p exhibited the greatest positive effect value and were strongly correlated (Figure 4). The strong correlation between miR-499a-5p and miR-361-5p implies that they may be functionally related or co-regulated in the context of ASD. Additionally, we observed a strong positive correlation between miR-150-5p and miR-3613-3p. It suggests that these two miRNAs may have also a cooperative or synergistic effect in influencing the development and manifestation of ASD. Based on these correlations, we selected one of the miRNAs that are strongly correlated with each other to be used for diagnostic purposes. These results may allow for the selection of a smaller subset of miRNAs that are most informative for ASD diagnosis. After highly correlated feature elimination, miR-3613-3p, miR-19a-3p and miR-126-3p were selected. As a second step of feature selection, Lasso regression analysis was performed to determine which miRNAs are the major ASD-associated miRNAs that have a significant impact on the detection of ASD. Which miRNA/miRNAs have a high impact on ASD detection?

After selecting miR-126-3p as the most promising miRNA for ASD detection, we trained the ML models on the training data. Five different algorithms were used: KNN, Logistic regression, SVM, RF and Decision tree and their results were presented in Table 2. The ML models have undergone training and are subsequently assessed using k-fold cross-validation. This approach is employed to gauge the accuracy and generalizability of the models. Multiple metrics are available to assess the models’ performance, including accuracy, precision, recall, F1 score, and others. In our study, we primarily reported the accuracy score as the key metric of interest.

**Table 2.** Model performance of machine learning classifiers on the validation set. The table summarizes the results obtained for each classifier, including accuracy, SD, specificity and sensitivity.

| **Models** | **Accuracy** | **SD** | **Specificity** | **Sensitivity** |
| --- | --- | --- | --- | --- |
| **KNN** | 0.94 | 0.06 | 0.81 | 1 |
| **Logistic Regression** | 0.94 | 0.06 | 0.81 | 1 |
| **Support vector machine** | 0.94 | 0.05 | 0.82 | 1 |
| **Random forest classifier** | 0.94 | 0.05 | 0.84 | 0.98 |
| **Decision Tree** | 0.96 | 0.05 | 0.9 | 0.99 |

The analysis of various machine learning models yielded accuracy scores exceeding 90% across the board. Notably, the K-nearest neighbors (KNN) model exhibited an impressive accuracy of 0.94, indicative of its proficiency in generating accurate predictions. Furthermore, the Logistic Regression model demonstrated a perfect sensitivity score of 1, signifying its exceptional ability to correctly identify positive instances, while achieving a specificity of 0.81, underscoring its precision in recognizing negative instances.

The Support Vector Machine (SVM) model showcased excellent performance, boasting an accuracy of 0.94, coupled with a high sensitivity of 1 and a perfect specificity of 0.82. The Random Forest (RF) classifier also achieved commendable accuracy at 0.94, complemented by a sensitivity of 0.98 and a perfect specificity of 0.84.

The top-performing model, however, emerged as the Decision Tree, with the highest accuracy score of 0.96. This exceptional accuracy was matched by an excellent sensitivity of 0.99, further bolstered by a commendable specificity of 0.9. These findings underscore the robust predictive capabilities of these machine learning models within the context of the study, with the RF model notably demonstrating superior overall performance.

The confusion matrix can also be seen in Figure 5.

**Figure 5.**
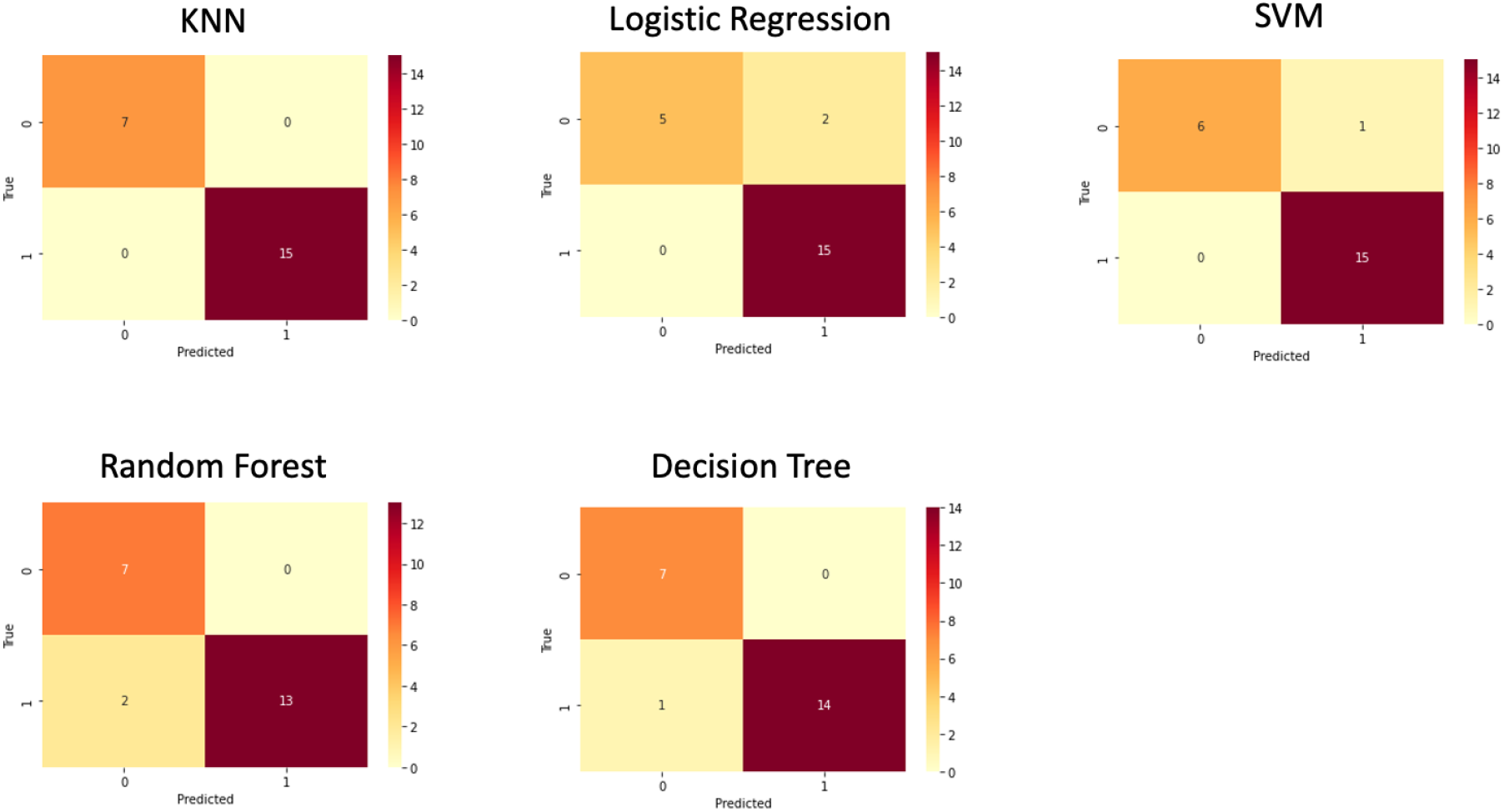
Confusion Matrix for all ML models

The confusion matrix was used to compare the different machine-learning algorithms.

To validate our models, we applied trained model on unseen test data set. KNN using miR-126-3p exhibited the best predictive performance among the five ML models (Table 3). The selected biomarker clearly discriminates healthy and ASD groups with over 90% accuracy for all ML models. These findings suggest that reduced levels of circulating miR-126-3p on the selected 6 miRNAs can serve as effective biomarker for ASD detection.

**Table 3.** Model performance of machine learning classifiers on the independent cohort. The table summarizes the results obtained for each classifier, including accuracy, SD, specificity and sensitivity.

| Models | Accuracy | Specificity | Sensitivity |
| --- | --- | --- | --- |
| KNN | 1 | 1 | 1 |
| Logistic Regression | 0.90 | 0.71 | 1 |
| Support vector machine | 0.95 | 0.85 | 1 |
| Random forest classifier | 0.90 | 1 | 0.86 |
| Decision Tree | 0.95 | 1 | 0.93 |

### 2.4. Functional Analysis

We conducted an extensive functional enrichment analysis of miR-126-3p, focusing on its involvement in specific molecular activities based on Gene Ontology (GO). The results, presented in Figure 6A, unveil noteworthy enrichments that provide insights into the regulatory capacity of miR-126-3p. Our comprehensive analysis demonstrated that miR-126-3p is intricately involved in 25 distinct pathways, highlighting its significant role in cellular regulation. Notably, one of the most prominent pathways in which miR-126-3p displayed high enrichment is the mTOR (mammalian target of rapamycin) signalling pathway, which governs critical cellular processes. Within the mTOR signalling pathway, miR-126-3p exhibited a substantial enrichment and was notably associated with three target genes: Insulin Receptor Substrate 1 (IRS1), Phosphoinositide-3-Kinase Regulatory Subunit 2 (PIK3R2), and Vascular Endothelial Growth Factor A (VEGF-A) (Figure 6B). This observation points toward a compelling regulatory role for miR-126-3p within the mTOR pathway, which, in turn, influences key cellular processes. IRS1 is a central mediator in insulin signalling, influencing cell growth and metabolism^20^. Its interaction with miR-126-3p in the context of the mTOR pathway suggests a regulatory link between miR-126-3p and metabolic processes. PIK3R2 is a regulatory subunit of the PI3K complex^21^, a key player in the mTOR pathway. The interaction between miR-126-3p and PIK3R2 suggests a potential role in modulating PI3K-mediated signalling. VEGF-A is a critical factor in angiogenesis, which is intricately connected with mTOR activity^21^. The association between miR-126-3p and VEGF-A hints at miR-126-3p’s potential role in regulating angiogenic processes.

**Figure 6.**
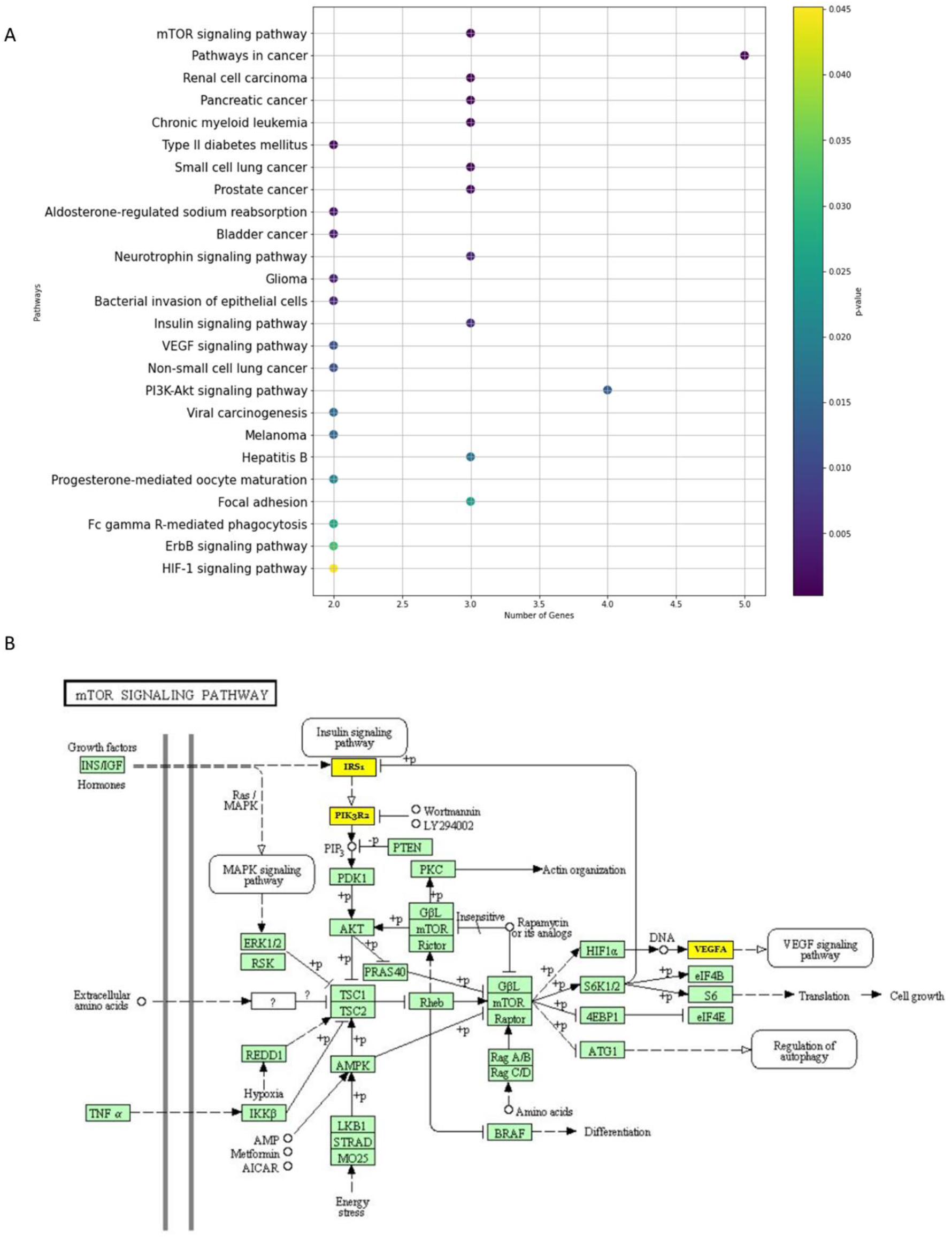
Functional enrichment analysis of target genes based on gene ontology regarding (A) molecular function (A) KEGG pathway analysis. B) mTOR pathway analysis IRS1, PIK3R2 and VEGFA

## 3. Discussion

miRNAs have attracted increasing attention in recent years as a crucial tool for gaining a deeper understanding of a wide range of biological processes, including the emergence of many human diseases such as cancer, cardiovascular problems, and neurological disorders field^22–24^. The quantitative measurement of small miRNAs could be established routinely in the clinical laboratory, but the interpretation of the results requires reliable and automatic ways. In our previous study, we investigated the miRNA expression profiles of the parents of autism patients, which provided important insights into the potential genetic and epigenetic factors underlying the development of ASD ^14^. The rationale behind conducting this study is to identify a reliable diagnostic biomarker panel for ASD using miRNAs. There is currently a need for a non-invasive, reliable, and cost-effective biomarker for ASD diagnosis, as current diagnostic methods are subjective and time-consuming.

These findings suggested that alterations in miRNA expression patterns may contribute to the risk of developing ASD, and further investigation of these miRNAs may be valuable in understanding the underlying mechanisms of ASD pathogenesis. Building on these previous findings, the current study aimed to identify specific miRNAs that could serve as diagnostic biomarkers for ASD, which is a challenging condition to diagnose accurately and early. The significance of the findings is that this study has identified a potential of miR-126-3p, which can serve as a diagnostic biomarker for ASD with high accuracy. This can potentially lead to earlier and more accurate diagnoses of ASD, allowing for earlier interventions and improved outcomes. Additionally, the use of ML, such as KNN, provides a promising method for accurately predicting ASD diagnosis using miRNA biomarkers.

We have demonstrated that our KNN model can accurately estimate the risk of ASD by using a dataset that contains miRNA levels. To our knowledge, this study uses a large dataset to develop the computer-aided diagnostic model. Our results show that KNN may have the potential to aid physicians in discriminating between healthy and patients with ASD.

Potential functional implications of miR-126-3p in molecular pathways gave offers valuable information for understanding the molecular interactions. We showed that miR-126-3p highly enriched in mTOR, a serine/threonine kinase, plays a pivotal role in regulating cellular responses to nutrient availability, influencing critical processes such as protein synthesis, cell growth, and proliferation^25–27^. Dysregulation in the mTOR pathway has been linked to the development of autism, contributing to aberrant synaptic protein synthesis and associated symptoms, including macrocephaly, seizures, and learning deficits^28–30^. Notably, approximately 8-10% of autism cases have been associated with abnormalities in the mTOR signaling pathway^30^. Moreover, a substantial proportion of autism predisposition genes directly or indirectly intersect with mTOR signaling activity^31^. These findings collectively underscore the central role of mTOR signaling in autism. Mainly it has been already showed that miR-126 directly targets the PIK3R2 gene^21^. The enrichment of miR-126-3p within the mTOR signaling pathway opens new avenues for research. It suggests that miR-126-3p’s regulatory role can significantly impact key cellular processes, such as cell growth, metabolism, and angiogenesis. Further in-depth investigations are warranted to unravel the precise molecular interactions between miR-126-3p and its target genes within the mTOR pathway.

It is worth noting that our study has some limitations. First, patients, since only one centre provided the results should be repeated with more samples. To confirm the efficacy of models and differentially expressed miRNAs, additional data using the same sample preparation processes and technic from other nations and ethnic groups are needed and future studies with larger sample sizes are needed to validate our results. Second, the sample size was relatively small, additionally, our study only focused on circulating miRNAs, and it remains to be determined if other types of miRNAs, such as tissue-specific miRNAs, can be used as diagnostic biomarkers for ASD.

Additionally, the strength of these findings is further emphasized by the fact that ASD is notoriously difficult to diagnose, with a lack of reliable and objective biomarkers. The identification of a panel of miRNAs that can accurately distinguish ASD from healthy subjects has important implications for early detection and intervention, ultimately improving outcomes for individuals with ASD.

In conclusion, our study provides evidence that miR-126-3p has high potential as a diagnostic biomarker panel for ASD. These findings have important implications for the early detection and treatment of ASD, which can improve outcomes for individuals with this disorder. Further studies are needed to validate these findings and to explore the underlying mechanisms by which these miRNAs contribute to the pathogenesis of ASD. In conclusion, our results imply that circulating miRNAs have the potential to serve as biomarkers for the identification of ASD. The miR-126-3p with all ML models showed the potential to be a useful diagnostic tool to advance the diagnosis of ASD.

## 4. Materials and Methods

### 4.1. Ethics Statement

The methods involving human participants in this study were conducted in accordance with the applicable guidelines and regulations set forth by the Ethics Committee of Erciyes University School of Medicine. The study received validation from the committee on September 20, 2011, with the assigned committee number: 2011/10. Furthermore, the study was approved by the Hospital. Prior to participating in the study, all parents provided written informed consent, which was obtained and validated by the Ethics Committee of Erciyes University School of Medicine (committee number: 09-20-2011, Ethics committee number: 2011/10). Patient selection criteria were also adhered to, and a comprehensive explanation of the study was provided to both the participants and their parents before their enrollment.

### 4.2. Data Set

The study included a total of 217 participants from both multiplex and simplex families. Among them, there were 45 subjects with ASD aged between 2 and 13 years (31 males and 14 females), along with 21 age- and sex-matched typical control subjects aged between 3 and 16 years (10 males and 11 females). Additionally, 33 healthy siblings aged between 1 and 20 years (17 males and 16 females) were included in the study. All participants were of Turkish origin.

For miRNA analysis, plasma samples were collected from patients and healthy controls, and miRNA PCR Array profiles were extracted, containing measurements for 372 miRNA expression levels.

Processing and analyzing of raw data were accomplished using R software (version 4.2.0, Limma package) to identifity differentially expressed miRNAs, adjusted p <0.05 were set as the threshold. Differential miRNA expression analysis were explained previously ^14^.

### 4.3. Data pre-processing – Feature Selection

The ML technique includes a crucial step called data pre-processing. Data selection, noise filtering, imputation of missing values, resampling (SMOTE) to handle imbalanced data, feature selection, and normalisation are all components of large-scale data preparation.

#### 4.3.1. Feature Selection

Log2-normalized microarray expression values extracted from serum samples served as the primary input for rigorous feature selection. The dataset was carefully partitioned into three distinct parts, with two out of the three parts allocated for the training dataset. Subsequently, a comprehensive analysis of the correlation matrix between miRNAs was executed using the Spearman’s correlation method exclusively within the training data set.

Following the identification of pertinent miRNAs, a robust repeated stratified k-fold validation framework was employed to rigorously apply the selection criteria. This entailed a partitioning scheme of three splits and repeating the process five times within the confines of the training set. This methodological approach was designed to circumvent the pitfalls of data overfitting and ensure the reliability of the results.

#### 4.3.2. Lasso

The LASSO (Least Absolute Shrinkage and Selection Operator) linear regression model was employed in this study to enhance prediction accuracy. The standard five-fold cross-validation technique assessed and validated the model’s performance. In recent years, LASSO regression analysis has gained prominence as a valuable tool for identifying diagnostic or prognostic features, and it has been widely utilized in various research studies ^32^.

Repeated 5-fold cross-validation was used split data into train and test sets using the Python (v3.8.8) and Python libraries “Sklearn”. Sklearn library was used for feature selection, Lasso, and Normalization/scaling for data.

Overfitting is prevented by using K-fold cross-validation. We also measure the model accuracy using 5-fold cross-validation. The data is equally divided into 5 folds (repeated 5 times) of which is used for training and 1 for evaluation. The cross-validation is carried out once again after rearranging the data set. Following training, only test data are used to evaluate the model’s prediction.

### 4.4. Machine Learning Algorithms

Recent research on the prediction and classification of disease using gene expression data has made extensive use of many algorithms created to solve classification problems in machine learning^33,34^. In machine learning, the general classification process entails training a classifier to detect patterns effectively from provided training samples and then classifying test data using the trained classifier. The classification process employs illustrative classification methods such as the k-nearest neighbour (KNN), support vector machine (SVM), random forest classifier and Decision Tree.

KNN, Logistic Regression, SVM, Random Forest, and Decision Tree were used to create diagnostic models for autism utilising five ML algorithms using Python (v3.8.8) and Python libraries “xgboost”, “TensorFlow”, “sklearn” and “imblearn”.

One of the most popular techniques for memory-based induction is the KNN. Based on similarity metrics, KNN retrieves the k closest vectors from the reference set from an input vector and uses the labels of the k closest neighbours to decide on the input vector’s label ^35^. Euclidean distance and Pearson’s coefficient correlation were utilised as the similarity metric.

Logistic regression is a vital statistical technique employed in various fields to model the relationship between binary outcomes and one or more predictor variables ^36^. This versatile method plays a crucial role in binary classification tasks. Logistic regression provides a way to estimate the probability of an event occurring based on the values of independent variables, making it a fundamental tool for decision-making and risk assessment^37^.

SVM is an effective technique for creating classifiers. To enable the prediction of labels from one or more feature vectors, it seeks to establish a decision boundary between two classes ^38^. The hyperplane, a decision boundary, is oriented to be as far away from each class’s nearest data point as is technically possible. These closest points are called support vectors.

An ensemble of learning techniques for classification, regression, and other tasks known as random forest (RF) is a machine learning algorithm that works by building a large number of decision trees during the training phase and then displaying the classes (for classification problems) or mean predictions (for regression problems) of the individual trees ^39^. The RF algorithm creates every decision tree it contains, training each one with a portion of the problem’s data. It randomly selects N records from the data, then uses multiple trees that were built and trained using the bagging principle to learn from them. This makes sure that every decision tree has a unique perspective on the issue. After all the decision trees have been trained, RF votes on all of them and then make decisions according to the classification or regression problem that needs to be solved ^40^.

Decision trees are recognized as a highly effective data mining methodology, widely embraced in diverse academic disciplines^41^. Their appeal is attributed to their user-friendly characteristics, interpretability, and robustness, even when confronted with missing data. It’s worth noting that decision trees demonstrate versatility by accommodating both discrete and continuous variables, which may function as either target or independent variables^42^.

Each ML model shares the same training and test sets. As shown in Figure 1, we train the machine learning models and measure training time at a training stage, then test their accuracy and measure prediction accuracy at a testing stage. To compare various ML methods and parameter settings, the model performance such as prediction accuracy and standard deviation were produced, as well as the area under the receiver operating characteristic curve.

### 4.5. Validation approaches

The performance of each machine learning algorithm underwent evaluation through two distinct forms of cross-validation. Initially, a random 5-fold cross-validation was executed by randomly assigning each patient to one of three groups. In each iteration of the cross-validation process, one group was set aside as the test set, while the classifiers were trained on the remaining data.

Subsequently, considering that the data originated from an independent cohort, the accuracy of all machine learning models was thoroughly assessed. Performance metrics, including sensitivity and specificity, were diligently evaluated to provide a comprehensive understanding of the algorithm’s effectiveness.

### 4.6. Pathway analysis and Gene Ontology (GO)

To gain a deeper insight into the functional significance of a chosen miRNA, a pathway prediction analysis was conducted. To accomplish this, the bioinformatics tool DIANA-mirPath (version 3) was employed^43^. This computational approach enabled the identification of all genes and cancer-related molecular pathways influenced by the selected miRNAs, providing valuable insights into their potential roles in biological processes.

### 4.7. Data availability statement

The data used in this study were collected by Ozkul and colleagues^14^. The research article titled ‘A heritable profile of six miRNAs in autistic patients and mouse models’ was published in Nature Scientific Reports in 2020 ^14^. The data set utilized in the study contains miRNA (microRNA) levels measured from human samples. Detailed information about the miRNA data set can be found in the supplementary materials of the article^14^.

### 4.8. Ethics Statement

All methods involving human participants were performed in accordance with the relevant guidelines and regulations by the Ethics Committee of Erciyes University School of Medicine and validated on <u>09-20-2011 by committee number: 2011/10</u>. This study was approved by the Hospital. All parents gave written informed consent before participation (Ethics Committee of Erciyes University School of Medicine 09-20-2011 committee number: 09-20-2011 Ethics committee number: 2011/10).

This work was made possible by a grant to Y. Ozkul from Tübitak, 1010 (EVRENA project ID 112S570), and to Minoo Rassoulzadegan Fondation Nestlé Rassoulzadegan 2019-2020.

## Author Contributions

Conceptualization, E.M., A.D., M.R., S.T. and Y.O.; methodology, E.M.., A.D., and M.R..; formal analysis, E.M.. and A.D..; writing—original draft preparation, E.M. and M.R..; writing—review and editing, A.D., S.T., Y.O. and M.R..; funding acquisition, Y.O and M.R..

## Competing interests

The author(s) declare no competing interests.

## Notes

https://www.nature.com/articles/s41598-020-65847-8#Sec34

### Competing Interest Statement

The authors have declared no competing interest.

